# Modeling Joint Abundance of Multiple Species Using Dirichlet Process Random Effects

**DOI:** 10.1101/056150

**Authors:** Devin S. Johnson, Elizabeth H. Sinclair

## Abstract

We present a method for modeling multiple species distributions simultaneously using Dirichlet Process random effects to cluster species into guilds. Guilds are ecological groups of species that behave or react similarly to some environmental conditions. By modeling latent guild structure, we capture the cross-correlations in abundance or occurrence of species over surveys. In addition, ecological information about the community structure is obtained as a byproduct of the model. By clustering species into similar functional groups, prediction uncertainty of community structure at additional sites is reduced over treating each species separately. The method is illustrated with a small simulation demonstration, as well as an analysis of a mesopelagic fish survey from the eastern Bering Sea near Alaska. The simulation data analysis shows that guild membership can be extracted as the differences between groups become larger and if guild differences are small the model naturally collapses all the species into a small number of guilds which increases predictive efficiency by reducing the number of parameters to that which is supported by the data.

## 1 INTRODUCTION

In recent years there has been considerable development of methodology for modeling and predicting abundance and occurrence of species of interest. Much of this development uses a hierarchical framework for developing models to fit the complexities of the observed data or natural abundance processes (Cressie *et al.*, 2009; Royle and Dorazio, 2008; Hobbs and Hooten, 2015). Some of these complexities may include: spatial and temporal dependence (Carroll *et al.*, 2010; Latimer *et al.*, 2009; Johnson *et al.*, 2013b; Thorson *et al.*, 2015; Ward *et al.*, 2010; Thorson *et al.*, 2016), nondetection of individuals at sampled sites (Dorazio and Connor, 2014; Royle, 2004), and zero-inflation (Johnson and Fritz, 2014; Thorson *et al.*, 2016). Many of these species distribution models (SDMs) were used to make inference to a single species or one-at-a-time modeling if community inference was desired. However, by not recognizing the fact that species interact, use of single species models for making inference for community abundance and structure can produce misleading results (Clark *et al.*, 2014). Hence, new joint species distribution models (JSDMs), which explicitly model species interactions (or, cross-correlation) have recently been developed (e.g., Dorazio and Connor, 2014; Latimer *et al.*, 2009; Thorson *et al.*, 2015, 2016). Herein, we propose a novel JSDM approach which models species interactions through membership in a latent ecological guild (Simberloff and Dayan, 1991) or functional group within the sampled range of habitats.

Typically, description of an abundance model begins with a generalized linear model (GLM) structure for the abundance process using a discrete value distribution such as Poisson or negative-binomial. For example, one might model the abundance as a Poisson observation with log-mean that is a function of covariates. Those covariates might include habitat variables or variables related to the sampling procedure which are thought to be related to the observed abundance. Alternatively, one might log transform the abundance and use Gaussian linear models (Johnson *et al.*, 2013b; Johnson and Fritz, 2014; Ward *et al.*, 2010), but the general mean structure is usually the same. Herein, we will focus on the GLM versions. The focus of the abundance modeling is related to either establishing an ecological relationship between (joint) abundance and the environmental covariates or predicting abundance at unsampled locations.

To extend the single species GLM oriented model to account for interactions of multiple species and improve prediction and inference of community structure and joint abundance, there have been several approaches which differ in the details of interaction modeling, but all were placed in the GLM framework by adding random effects which are either directly correlated between species (Clark *et al.*, 2014; Dorazio and Connor, 2014; Latimer *et al.,* 2009) or when marginalized from the (log-linear) model imply a cross-species correlation structure (Thorson *et al.,* 2015, 2016). The direct approach of using a free parameter for every pair of species when modeling the species-level correlation has been successfully implemented (Clark *et al.*, 2014; Latimer *et al.*, 2009), however, in those studies there were a high number of sampled sites or a low number of species considered. In other studies, unstructured covariance did not produce reliable results (Dorazio and Connor, 2014). Thus, recent efforts to contribute novel methodology for JSDMs have focused on reducing the number of parameters used to model species interactions. Dorazio and Connor (2014) used a known species trait proximity matrix to model the species-level covariance matrix using a spatial correlation function. By using the known information on species similarity there are only two parameters necessary to model the cross-correlation. Another low complexity approach has been proposed (Thorson *et al.*, 2015, 2016) using linear combinations of latent random effects. Specifically, the latent effects are spatial fields, but the same methodology could be applied using independent random effects. If the number of random effects is small relative to the number of species modeled, the number of parameters necessary for modeling species cross-correlation can be significantly reduced from the unstructured scenario.

As a novel alternative, we propose a JSDM that uses latent ecological guilds to model interactions among species and obtain joint abundance inference. Herein, we also consider joint species occurrence as well, where occurrence is defined as the binary presence (i.e., abundance > 0) or absence (abundance = 0) of a species. Dorazio and Connor (2014) used known guild membership of different species to model independence of some species in a crosscorrelated JSDM. Simberloff and Dayan (1991) defines an ecological guild to be “a group of species that exploit the same class of environmental resources in a similar way.” With this definition in mind, we seek to build a model where species are cross-correlated in abundance because there are unknown group effects for some set of covariates, i.e., if the group (guild) structure was known they could be represented by (group × covariate) interaction terms in the abundance GLM models. To accomplish this task we format the model as a latent class or mixture model (see McLachlan and Peel, 2004). Mixture models or latent class models are often used to model dependance between variables in a nonparametric fashion because for a sufficiently large number of groups, marginalizing over the random latent classes can approximate any dependence structure to whatever degree desired (McLachlan and Peel, 2004; Vermunt *et al.,* 2008). It has been shown that this holds even when the conditional models are independent given group membership (Dunson and Xing, 2009). In an ecological abundance context, finite mixture models have been used in the past to model spatial heterogeneity and correlation in a nonparametric fashion (Dorazio *et al.,* 2008; Johnson *et al.,* 2013b). In this paper we take inspiration from nonparametric dependence methods used for spatial association and apply it to species interaction in abundance modeling.

In the following section we describe the general infinite mixture framework using latent groups and describe the Dirichlet Process (DP) for modeling group membership and the number of groups. There are several models choices for number and assignment of latent classes, but we utilize the DP due to its long history and good clustering properties (Casella *et al.,* 2014). Parameter estimation in the DP-JSDM is challenging due to the latent class process. We provide a reversible-jump MCMC (RJMCMC; Green 2003) algorithm for making Bayesian inference. Finally, we apply the method to few simulated data sets, as well as, a real data set on mesopelagic fish communities in the eastern Bering Sea, Alaska.

## 2 METHODS

### 2.1 General model framework

We begin the description of the proposed methods with some notation. First we assume there are *J* surveys, for which abundance (or count index; hereafter we use the term “counts”) of *I* different species is measured. Let *n_ij_* be the observed count for *i*th species in survey j. We also use the vector notation *n_i_* = (*n*_*i*1_,…, *n_iJ_*)′ and 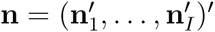 for the *n_ij_*, as well as, other quantities described later. For occurrence modeling we denote occurrence as *y_ij_* = 1 if *n_ij_* > 0 otherwise *y_ij_* = 0. In practice, *n_ij_* need not necessarily be observed for occurrence modeling. The notation y_i_ and y are similar to the abundance counterparts.

For abundance modeling, there are several possible distributions that could be used to model the observed discrete counts, Poisson, negative binomial, zero-inflated Poisson, etc., so we will generically denote this observation model as [*n_ij_*|*z_ij_*, γ] where *z_ij_* is a latent Gaussian variable controlling the level of expected abundance and γ is a set of, possibly nuisance, parameters. The notation “[*A*|*B*]” refers to the conditional distribution of *A* given *B*. For example, if a Poisson distribution is considered,

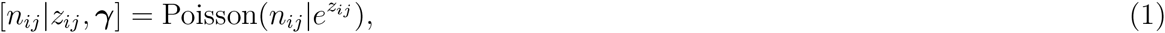

and γ is not necessary. In the example analysis of mesopelagic fish surveys we utilize a zero-inflated Poisson (ZIP) model, so,

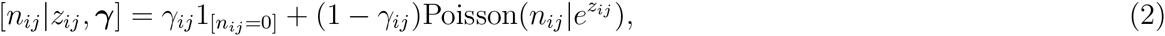

the additional γ*_ij_* parameter is the mixing probability for the extra zeros. For occurrence modeling we use

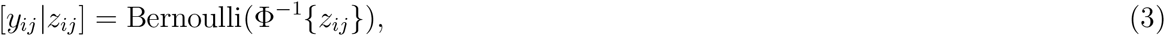

where Φ(·) is the standard normal CDF, that is, a probit link function.

To account for unknown interspecies correlations we take a clustering approach inspired by the analysis of Johnson *et al*. (2013b) for incorporating spatial structure when there are no reasonable distance metrics or neighborhood groupings are unknown. The model is constructed by envisioning an unknown partition, *p*, of the species into *κ_p_* groups such that species within groups behave similarly with respect to the abundance process. For a given *p*, we model (in vector form) the latent **z** process with the linear model

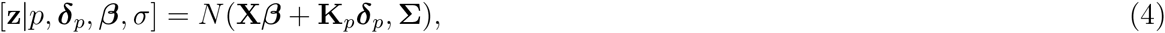

where

- **X** is a design matrix of covariates for which there are no group-level effects,
- *β* is a vector of regression coefficients,
- **K***_p_* = **C***_p_* ⊗ **H**, where **C***_p_* is an *I* × *κ_p_* binary matrix indicating which species belong to each group in *p* and **H** is a *J* × *q* matrix of q habitat covariates recorded at the *j*th survey,
- 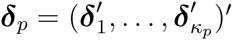 is a vector of normally distributed random effects, where, 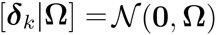, for *k* = 1,…,*κ_p_*
- Σ is a diagonal matrix with entries 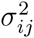 (for occurance modeling *σ_ij_* = 1).

To reduce the complexity of the proposed model we suggest the following for general practice:

i. for abundance models, set *σ* = diag(Σ^1/2^) = exp{**L*θ***}, where **L** is a matrix of design covariates and
ii. set **Ω** = *ω*^2^(**H′H**)^−1^, where *ω* = exp(*ξ*).

With respect to (*i*), there are some useful special cases, namely, **L** = **1** gives *σ_ij_* = *σ* and **L** = **I**_*I*_ ⊗ **1**_*J*_ gives *σ_ij_* = *σ_i_*. However, the overdispersion parameters could also be modeled based on covariates associated with sampling methods, etc. Suggestion (*ii*) was formulated from the covariances of the *g*-prior (Tiao and Zellner, 1964). The *g*-prior, *N*(**0**, *ω*^2^(**H′H**)^−1^), is an often used prior for regression coefficient parameters. It has the nice benefit that, with a single parameter, it automatically controls the scale of variance and covariance for each coefficient based on the scale of the covariates and their cross-correlation. The exponential reparameterization is used for ease of inference, that is *ξ* can be unconstrained.

The previous description assumed that the correct partitioning of the species is known, however, for most real data sets, the correct partition is unknown. Thus, we must also provide a probability model over partitions, [*p*|*α*], such that marginalization over the unknown partitions creates random coefficient vectors that are nonparametric in their distribution. A commonly used distribution over partitions is the Chinese Restaurant Process (CRP). A construction definition of the CRP is described as follows, for a given parameter *α* > 0,

1. A customer enters the restaurant and sits at one of an infinite number of tables.
2. The next customer enters and chooses to sit at the occupied table with probability 1/(1 + *α*) or a new table with probability *α*/(1 + *α*).
3. In general,the *i* + 1 customer sits at an occupied table with probability proportional to the number of customers already seated or chooses an unoccupied table with probability proportional to *α*.

Under the CRP model individuals are exchangeable, i.e., individuals join groups based only on how many other individuals are in the group, not who else is in the group. This fact forms the basis for Bayesian inference for the CRP model via MCMC (Neal, 2000). The density function for the CRP cluster model is given by,

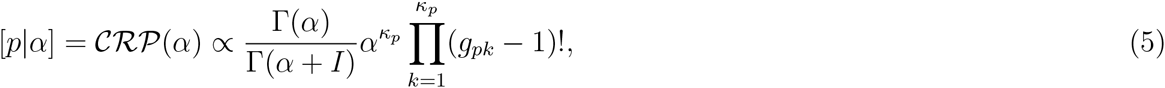

where *g_pk_* is the size of the *k*th group in *p*. Note, that the distribution of *p* is only a function of the number and sizes of the groups. Realizations of *p* with the same number of groups and groups sizes have the same probability regardless of which individuals fall in which cluster.

The Dirichlet process is connected to the CRP process because a DP process is constructed using the same procedure to seat the guests in the CRP model. Specifically, in terms of (4), let 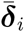 be the coefficient associated with the ith species, that is 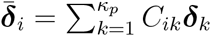, where *C_ik_* is the (*i*, *k*) entry of the **C***_p_* matrix. Now, if 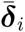 follows a DP then, conditionally,

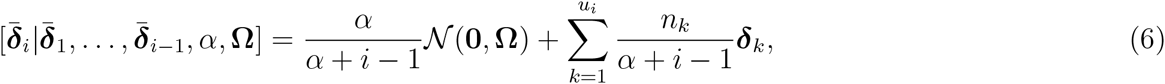

where *u_i_* is the number of unique values, ***δ**_k_*, of 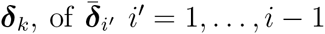, and *n_k_* is the number of species 1 through *i* − 1 belonging to group *k*. In other words, a new table is represented by a new value of ***δ**_k_*. Thus, the CRP partitioning combined with the ***δ*** realizations for each group implies that 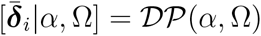.

Like the spatial covariance model use by Dorazio and Connor (2014), the DP-JSDM also marginally possesses generally positive cross-covariance structure. This makes intuitive sense as one is grouping similar species together or, if species are dissimilar, allowing them to be independent. The covariance structure of the DP-JSDM can be derived by forming an intercept random effect, ***η*** = **K***_p_**δ**_p_*, such that **z** = **X*β*** + ***η*** + ***ϵ***, where [***ϵ***] = *N*(**0**, Σ). Then, conditioning on the cluster assignment, the covariance matrix of the random effect ***η*** is,

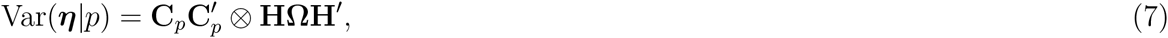

and the marginal variance is given by the mixture,

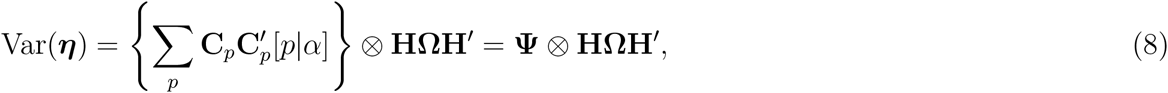

where **Ψ** is a matrix with (*i*, *i′*) entries equal to the probabilities that species i shares a guild with species *i′*. We term the **Ψ** matrix to be the species proximity matrix due to the fact that it forms a distance, of sorts, in the guild space of the species. Although, the covariance is never negative between any two species, it can be zero, thus those species that occupy different guilds will have uncorrelated *η* random effects, i.e., if *ψ_ii′_* ≈ 0, then Cov(*η_ij_*, *η_i′j_*) ≈ 0.

It should be noted, however, that although the covariance of the ***η*** random effect is generally, positive, that does not mean that there are only ‘positive’ (or zero) relationships between species. The clustering is based on the relationship each species has with the chosen covariates. For example, one species may react positively along a covariate gradient (*δ_i_* > 0) and another reacts negatively along that same gradient (*δ_i′_* < 0), therefore if a new site has a high level of this covariate, the first species will be predicted to be relatively abundant, while the other species abundance will be lower.

### 2.2. Bayesian inference

Because of the hierarchical and variable dimensional nature of the parameter space of the DP-JSDM model we employ a Bayesian approach via MCMC (Markov Chain Monte Carlo) for model fitting and inference. The posterior distribution of interest is given by

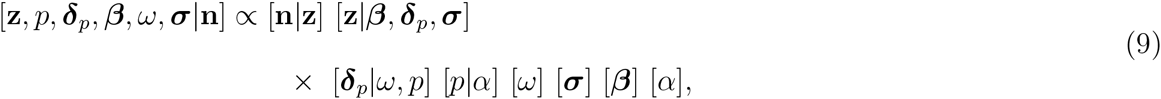

where [*ω*], [***σ***], [*β*], and [*α*] are the prior distributions for the parameters.

There are several derived parameters which may be of interest for making desired ecological inference. First, are predictions of community abundance at new locations or times. Second, one may be interested in the overall effect of the environmental covariates for a particular species represented by 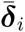. Finally, the matrix 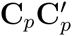 is an *I* × *I* indicator that a species is in the same guild (associated with) another species. The posterior mean of 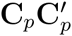 provides estimated guild proximity matrix, **Ψ**. Finally, the number of guilds, *κ_p_* (number of columns in **C***_p_*) may be of interest.

The most direct way to make inferences on the proposed hierarchical clustering model is through a reversible-jump Markov chain Monte Carlo (RJMCMC) algorithm (Green, 2003) to sample the posterior distribution of the parameters and clustering assignment. Here, we provide an overview of the RJMCMC, additional details of the sampler are given in Supplementary Material A.

In our description, we will assume the following prior distributions for the parameters:

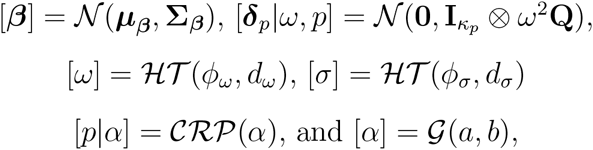

where **I***_κp_* is an identity matrix of size *κ_p_*, **Q** is a known positive-definite matrix, 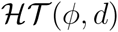 represents a scaled half-t distribution with scale parameter *ϕ* and *d* degrees of freedom, and 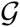 represents a gamma distribution with parameters *a* and *b*. For most of these parameters, the priors can be adjusted to whatever distribution the user would like, the trade-off being a Metropolis-Hastings (MH) update instead of a Gibbs step (e.g., for *β*) or no difference at all if the parameter has to be updated with an MH step to begin with (*ω*, *σ*, and *β*). However, the normal [***δ**_p_*|*ω*, *p*] prior is necessary to the proposed RJMCMC algorithm. Although, the known **Q** is not necessary. This is not as critical as it sounds as the marginal distribution is still a nonparametric DP process we just require that the base distribution be a multivariate normal.

The majority of the proposed RJMCMC algorithm is a standard Metropolis-within-Gibbs (hybrid) sampler for a GLM-like model. Conditioned on a realization of p, all the other parameters can be updated with a traditional MH step or a Gibbs step. Hence, we do not focus on their updates here (see Supplementary Material A). However, to update p, the dimension of the ***δ**_p_* vector will potentially change, necessitating the trans-dimensional aspect of the RJMCMC. Naively, the trans-dimensional moves require a joint (*p*, ***δ**_p_*) proposal which can be rejected often if one of those quantities is a bad fit for the current state of the remaining parameters even though the other is acceptable. Second, proposing new p such that the MCMC chain will mix well over the space of partitions is itself challenging. Because we are assuming that [**z**|*β*, ***δ**_p_*, ***σ***] and [**δ***_p_*|ω,p] are multivariate normal, the first problem can be handled with the partial-analytic RJMCMC method proposed by Godsill (2001) and utilized by Johnson and Hoeting (2011) and Johnson *et al.* (2013b) in similar trans-dimensional MCMC applications. The partial-analytic method allows proposal of a new model (*p* in this case) without jointly proposing the associated model specific parameters (δ_p_) because they can be analytically marginalized. This is a special case of a collapsed Gibbs sampler (Van Dyk and Park, 2008).

To produce efficient moves through guild space we use the the “individual links” definition of the the CRP process proposed by Blei and Frazier (2011) and subsequently used by Johnson *et al.* (2013b) for clustering spatial abundance trends. The links version of the CRP process is constructed as follows:

1. A customer enters the restaurant and sits at one of an infinite number of tables.
2. The next customer enters and chooses to sit with the first customer with probability 1/(1 + *α*) or a new table with probability α/(1 + *α*).
3. In general, upon entering the restaurant, the *i* + 1 customer sits with a previous *customer* (not a table) with probability proportional to 1 or the new customer sits by himself (self-links) with probability proportional to *α*.
4. Groups are constructed by collecting all cliques of the mathematical graph formed by the links between customers.

Blei and Frazier (2011) show that this definition of the CRP process is equivalent to the traditional definition presented previously. However, MCMC sampling is now based on sampling independent links between individuals. In terms of the multiple species model, let *ℓ_i_* ∈ {1,…, *I*} be the link for the ith species. The full conditional distribution of *ℓ_i_* is,

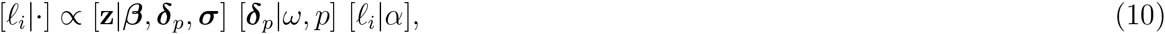

where *p* is the partition constructed from all *ℓ_i_* and,

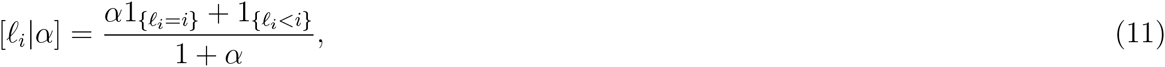

and 1_{·}_ is an indicator function for the condition in the brackets. It would be tempting to sample from this discrete distribution in Gibbs fashion, however, note that it depends on ***δ**_p_* which may be of different dimension under a different value of *ℓ_i_*. We can collapse over ***δ**_p_* and use the marginal distribution

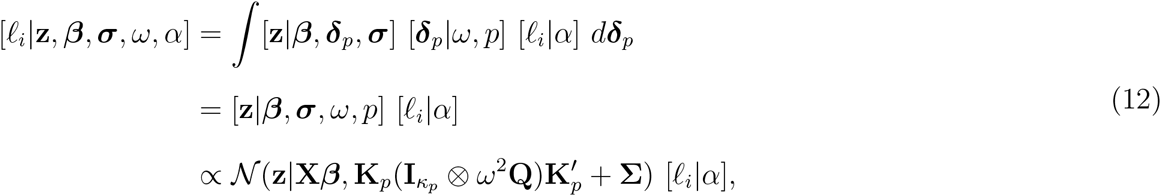

which does not depend on ***δ**_p_*. This approach was used by Johnson and Hoeting (2011) and Johnson *et al.* (2013b) exactly as described, however, we found that for a large number of species and samples, the covariance matrix 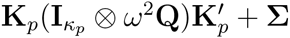 may be quite large and the inversion necessary to evaluate the [*ℓ_i_*|**z**, *β*, ***σ***, *ω*, *α*] for each species and potential link would make the chain prohibitively slow in practice. So, we sought an alternative formulation of the marginal distribution that did not require inversion of such a large covariance matrix. Using Laplace’s method (see Kass and Raftery 1995, Section 4.1) we can write

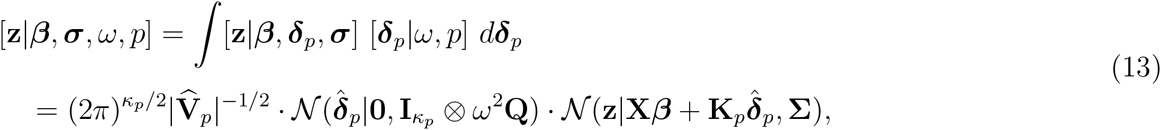

where 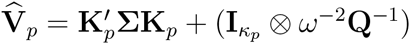 and 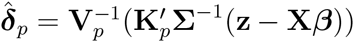, which are respectively the inverse covariance and mean for the Gaussian full conditional distribution [***δ***_p_|**z**, *β*, ***σ***, *ω, p*]. This is the same distribution used to update ***δ***_*p*_ with a Gibbs step following an update of p. Normally, Laplace’s method produces an approximation to the integral, but in this case the approximation is exact because the log integrand is quadratic in ***δ***_*p*_ (Goutis and Casella, 1999). By writing the integral in this way we need only invert Σ, which is diagonal, and **Q** because (**I***_κp_* ⊗ *ω*^2^Q)^−1^ = **I***_κp_* ⊗ *ω*^−2^**Q**^−1^. If we use **Q** = (**H′H**)^−1^ as previously suggested, then the inverse is trivial. Because, [*ℓ_i_*|**z**, *β*, ***σ***, *ω, α*] is relatively cheap to evaluate for each *ℓ_i_* we can use a Gibbs step and draw from the discrete distribution [*ℓ_i_*|**z**, *β*, ***σ***, *ω, α*] for each *i* = 1,…, *I*, with [**z**|*β*, ***σ***, *ω, p*] evaluated using (13) instead of (12).

## 3 A SIMULATION PROOF-OF-CONCEPT

To examine the ability of the DP-JSDM model to make inference to species interaction, as well as, to make community abundance predictions, we tested the model and RJMCMC sampler with a small group of simulated data sets. In analyzing the simulated data our objective was to assess whether the DP-JDSM model would, in practice, produce generally correct estimates of the guild structure. Second, would the DP-JSDM exhibit the expected behavior that as *ω* becomes small, the number of guilds (groups) estimated will go to one as the functional differences between the guilds (with respect to the variables in **H**) becomes insignificant.

### 3.1 Simulation and Analysis

Data were simulated for *I* = 20 species, *J* = 35 samples, and *κ_p_* = 5 groups. Six data sets were simulated corresponding to *ω* equal to 0.25, 0.5, 0.75, 1, 1.5, and 2. While the true number of groups is always technically equal to five, the practical differences between the groups tends to zero as ω becomes smaller. The group sizes were *g_pk_* = 7, 5, 4, 3, and 1. Three environmental variables composing the guild design matrix **H** were generated from a standard normal distribution. In addition, a single survey effort variable, **x** was generated to adjust overall abundance measurement. The global design matrix was set to **X** = [**1**, **x**, **H***_x_*], where **H***_x_* = [**H′**| … |**H′**]′, that is, **H** matrix is concatenated *I* times over species. Thus, ***δ**_p_* denotes guild differences from the overall global effect of the environmental variables, **H**. In order to maintain identifiability, we imposed the constraint that 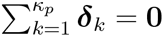. The global coefficient was set to *β* = (2, 1, 0, −1, 0.5)′ and each ***δ**_k_*; *k* = 1,…, 5, was drawn from *N*(**0**, *ω*^2^**H′H**). In these simulations all *σ_ij_* = 0, therefore, **z** ≡ **X*β*** + **K***_p_**δ**_p_*. However, a common *σ* was estimated in each analysis using a Poisson observation model, that is, [*n_ij_*|*z_ij_*] = Poisson(*e^z_ij_^*).

The prior distributions used were the same as specified in Section 2.2, specifically,

- 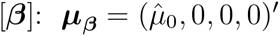 and 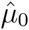 is the log of the mean observed count and Σ*_β_* = 100(**X′X**)^−1^.
- [*ω*]: *ϕ_ω_* = 1 and *d_ω_* = 1 which implies a half-Cauchy prior distribution.
- [*σ*]: *ϕ_σ_* = 1 and *d_σ_* → ∞ which implies a half-normal prior distribution.
- [*α*]: *a* = 0.258 and *b* = 0.038.

The prior distribution parameters for the gamma distribution [*α*] were chosen based upon the method of Dorazio (2009) with one alteration. Dorazio (2009) used the method to choose *a* and *b* such that the prior distribution over the number of groups was approximately uniform, that is, [*κ_p_*] ≈ 1/*I*, *κ_p_* = 1,…, *I*. However, we agree with the philosophy of Casella *et al.* (2014) that *a priori* we should prefer fewer groups, therefore, using the same optimization approach as Dorazio (2009), we chose a and b such that, approximately, [*κ_p_*] ∝ 1/*κ_p_*. So, all else being equal, a smaller number of groups is *a priori* preferred.

For each of the six simulated datasets, we sampled the posterior distribution (9) using the RJMCMC algorithm detailed in Supplementary Material A. Each sample consisted of 50,000 iterations following a burnin of 10,000 iterations. We created the `multAbund`^†^ package for the `R` statistical environment (R Development Core Team, 2015) which contains the code to run the RJMCMC algorithm described in Supplementary Material A.

### 3.2. Simulation results

As expected, when *ω* became small the DP-JSDM model was not able to distinguish guild differences between the species and essentially estimated one single group (Figure 1; *ω* = 0.25).

**Figure 1.**
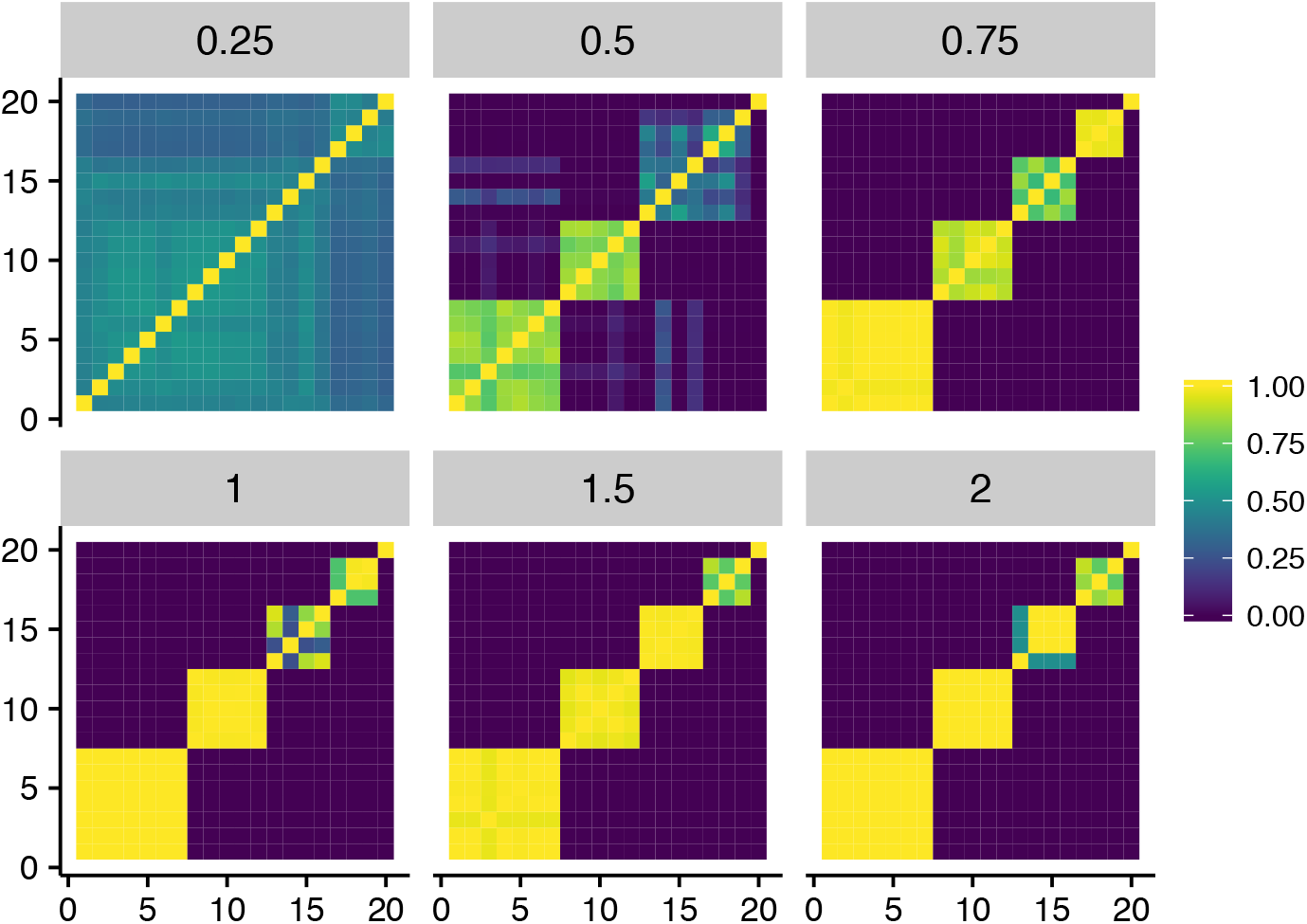
Estimated probabilities of joint guild membership between each species. For each panel, the value of *ω* used to simulated the data is provided in the bar above the plot.

As *ω* increased and guild differences became apparent the model was able to separate the species into their respective guilds reasonably well (Figure 1). In addition, as *ω* became large the precision with which the number of guilds was estimated increased as well (Figure 2).

**Figure 2.**
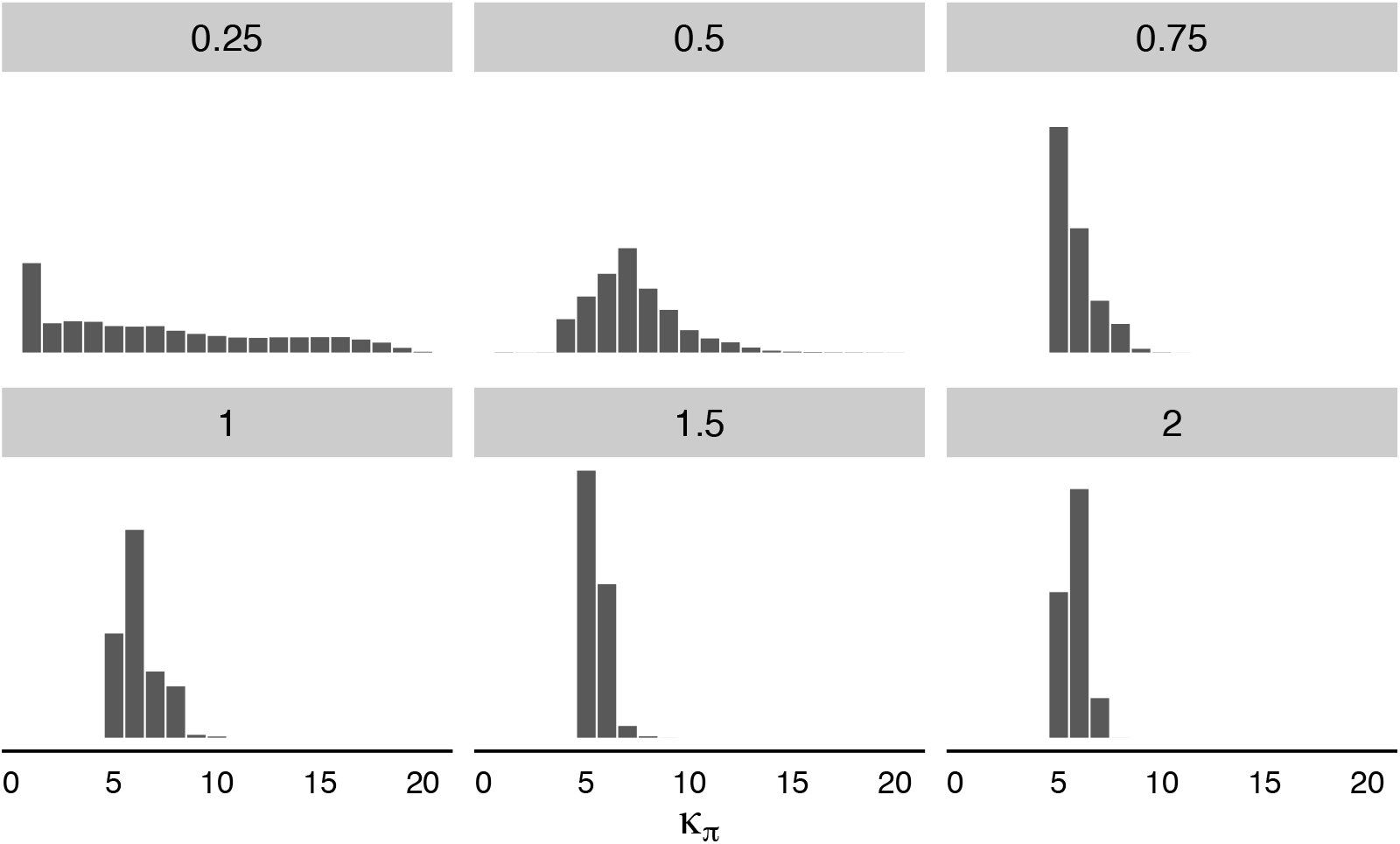
Estimated number of guilds, *κ_p_*, for simulated Poisson data sets with *ω* ranging from 0.25 to 2. For each panel, the value of *ω* used to simulate the data is provided in the bar above the plot.

There may be some bias as a few of the simulation runs produced K_p_ = 6 (Figure 2; *ω* =1 and 2), however, a full simulation experiment would be necessary to assess that fact. Even though we strived to create an efficient RJMCMC algorithm, it is still somewhat computationally intensive.

## 4 EXAMPLE: MESOPELAGIC FISH ABUNDANCE

### 4.1. Data

In our next demonstration of the DP-JSDM we analyze community structure and abundance of fishes that migrate diurnally between three mesopelagic depths in the eastern Bering Sea near Alaska. The tendency for most mesopelagic species to vertically migrate makes them an important trophic link between the deep scattering layer and upper surface waters (Sinclair *et al.,* 2015) yet, fundamental aspects of multi-species distributions and relative abundances have not been previously described.

The field effort identified three primary sample stations over highly productive areas of the eastern Bering Sea pelagic (Figure 3).

**Figure 3.**
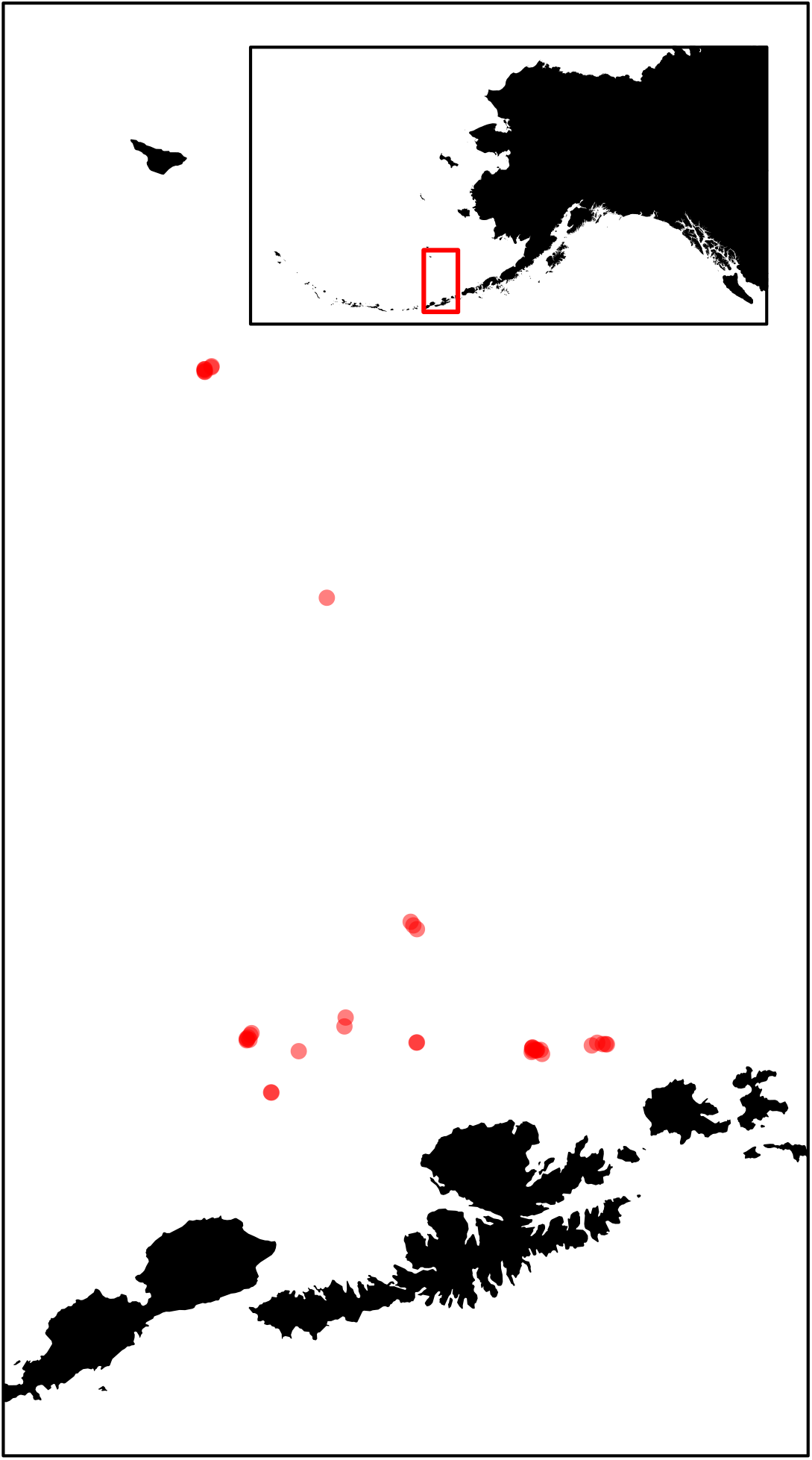
Locations of the mesopelagic trawl surveys. There were *J* = 41 separate trawl surveys used in the analysis of Section 4, however, some surveys were attempted geographically near other surveys, so, they are somewhat obscured in the figure.

In the summers of 1999 and 2000 a total of 29 daytime and 16 nighttime trawls were conducted at three depths (250, 500, and 1000 m) during a narrow sampling period. Four of these trawls were not analyzed due to technical difficulties in the field and we discarded them, resulting in *J* = 41 samples. Trawls were run at-depth for 15–90 minutes resulting in collections of over 50,000 individuals representing 55 species of fish and squid. Essentially, each individual trawl sample represents a community as influenced by depth and time of day. Here we will demonstrate the DP-JSDM using *I* = 20 of the relatively most common fish species (as opposed to squids, etc.). Many of the species were extremely rare in the survey effort (i.e., one individual observed over the entire study) and were removed.

The variables we put in the **H** design matrix reflect the belief that the species segregate into guilds based on diurnal vertical migration characteristics. So, the guild covariates recorded for each trawl are daylight cycle (day or night) and depth category (250, 500, or 1000 m). Here we used the full interaction model to define the **H** design matrix (i.e., `‘~ cycle*depth’` in `R` language model syntax). Because the duration of the trawl varied from survey to survey, the duration was included in the **X** matrix to model the overall abundance of fish caught in the trawl.

### 4.2 Model and analysis

Initial attempts at fitting a DP-JSDM proceeded in the same manner as the analysis of the simulation data in the previous section. Namely, we used the same Poisson model for the observed abundance counts. However, after initial fittings it became evident that the trawl data set possessed a significant level of zero-inflation relative to the Poisson distribution. This is likely due to the spatial patchiness of pelagic fish occurrence distributions (Benoit-Bird and Au, 2003). In addition, there may also be detection issues in the survey such that a zero count in the trawl does not necessarily mean absence of the species. However, unlike Dorazio and Connor (2014) we do not have replicated surveys at the same site and time in which to separate detection and absence. Therefore, we utilized a zero-inflated Poisson (ZIP) model in place of a Poisson GLM. The ZIP model used for this analysis is

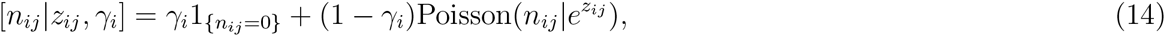

where 1_{*n_ij_*=0}_ is an indicator of a zero count and *γ_i_* is a species-specific zero-inflation mixture. We used the prior distribution,

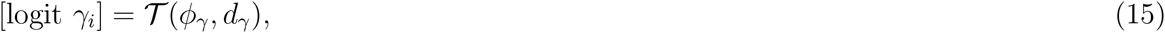

with scale parameter *ϕ_γ_* = 1.5 and degrees of freedom *d_γ_* = 6. This t distributed prior implies a prior distribution for γ_i_ that is approximately uniform over (0,1). For the remaining parameters we used the same prior specification as the simulated data analysis of Section 3.1.

To assess if there is any improvement gained by using the DP-JSDM we also fitted the ‘independent species’ JSDM, that is *κ_p_* = *I*, to the data. The JSDM we fitted was did not truly treat each species independently because there are shared terms in the **X** design matrix (i.e., trawl duration) but it allows us to assess improvement in classifying animals into functional guilds relative to cycle and depth over treating them separately. To ascertain the magnitude of improvement we would have liked to be able to use the ‘leave one out’ Bayesian predictive information criterion (BPIC) given by

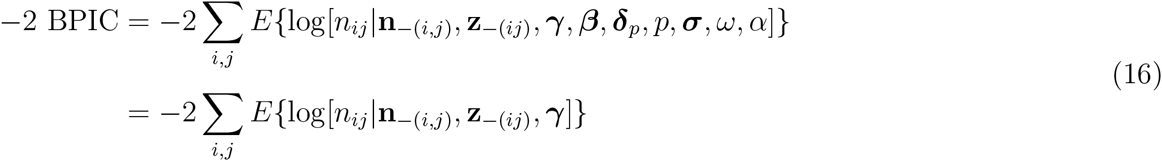

where **n**__(*ij*)_ is a vector of all observed data except *n_ij_* and log[*n_ij_*|**n**__(*i,j*)_, *γ*, *β*, ***δ**_p_*, *p*, ***σ***, *ω*, *α*] is the log posterior predictive density for the (*i*, *j*)th observation. However, it would be computationally infeasible to rerun the RJMCMC for every left out (*i*, *j*) entry. So, we used the ‘Widely Applicable Information Criterion’ (WAIC; Watanabe (2013)) as an approximation (Watanabe, 2010; Link and Sauer, 2016) to −2 BPIC, where

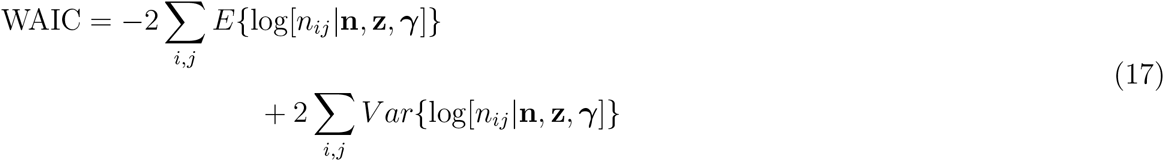

The WAIC requires only one run of the RJMCMC with the full data set. There are also other selection methods applicable, (Hooten and Hobbs, 2015), however, we found WAIC straightforward to implement for the DP-JSDM.

The model was fitted using the R package multAbund. The RJMCMC algorithm was run for 100,000 iterations following a burnin of 10,000 iterations. The package contains code to fit the Poisson abundance data model as well as the ZIP and Bernoulli probit model for occurrence. In addition to the joint analysis of abundance, we also analyzed the trawl survey data as an occurrence data set where *y_ij_* = 1 if *n_ij_* > 0, else *y_ij_* = 0. The occurrence analysis results are presented in Supplementary Material C.

### 4.3 Results

After fitting the ZIP version of the DP-JSDM and the independent species JSDM we noted there was a substantial improvement in WAIC under the DP-JSDM. WAIC for the DP-JSDM model was 3052.071 and WAIC = 3078.992 for the independence model. The posterior mode of the number of guilds was 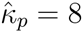 with 95% of the posterior probability mass falling on *κ_p_* = 8 or 9 guilds. Figure 4 depicts the estimated posterior matrix, 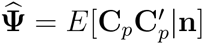 which defines the probability that any two species share the same vertical migration guild.

**Figure 4.**
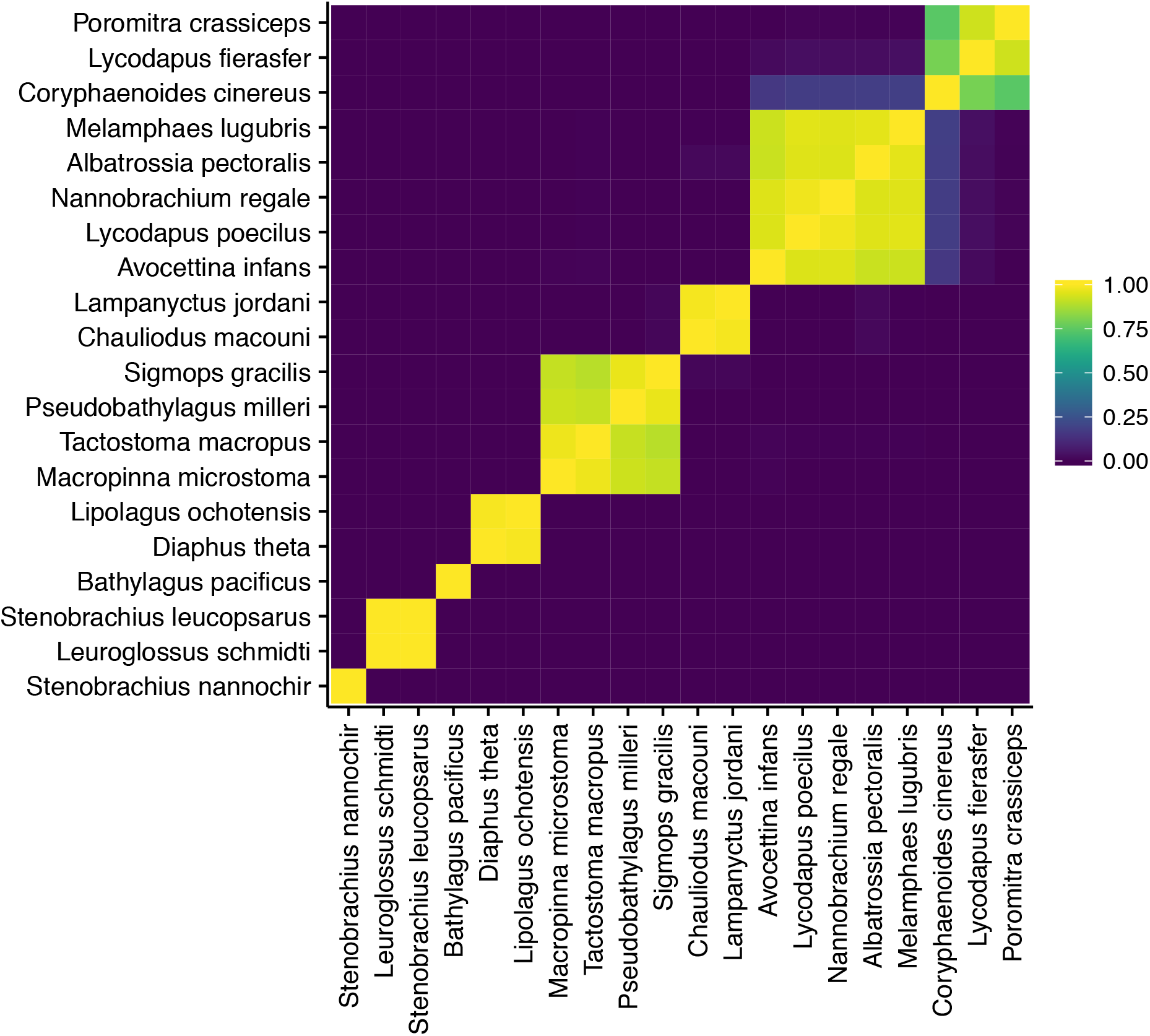
Estimated probability of joint guild membership for 20 of the fish species in the trawl survey with respect to abundance.

Using 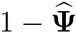 as a measure of distance between species, we plotted the species according to the associated dendogram (Figure 5), which gives a better visualization of the groupings.

**Figure 5.**
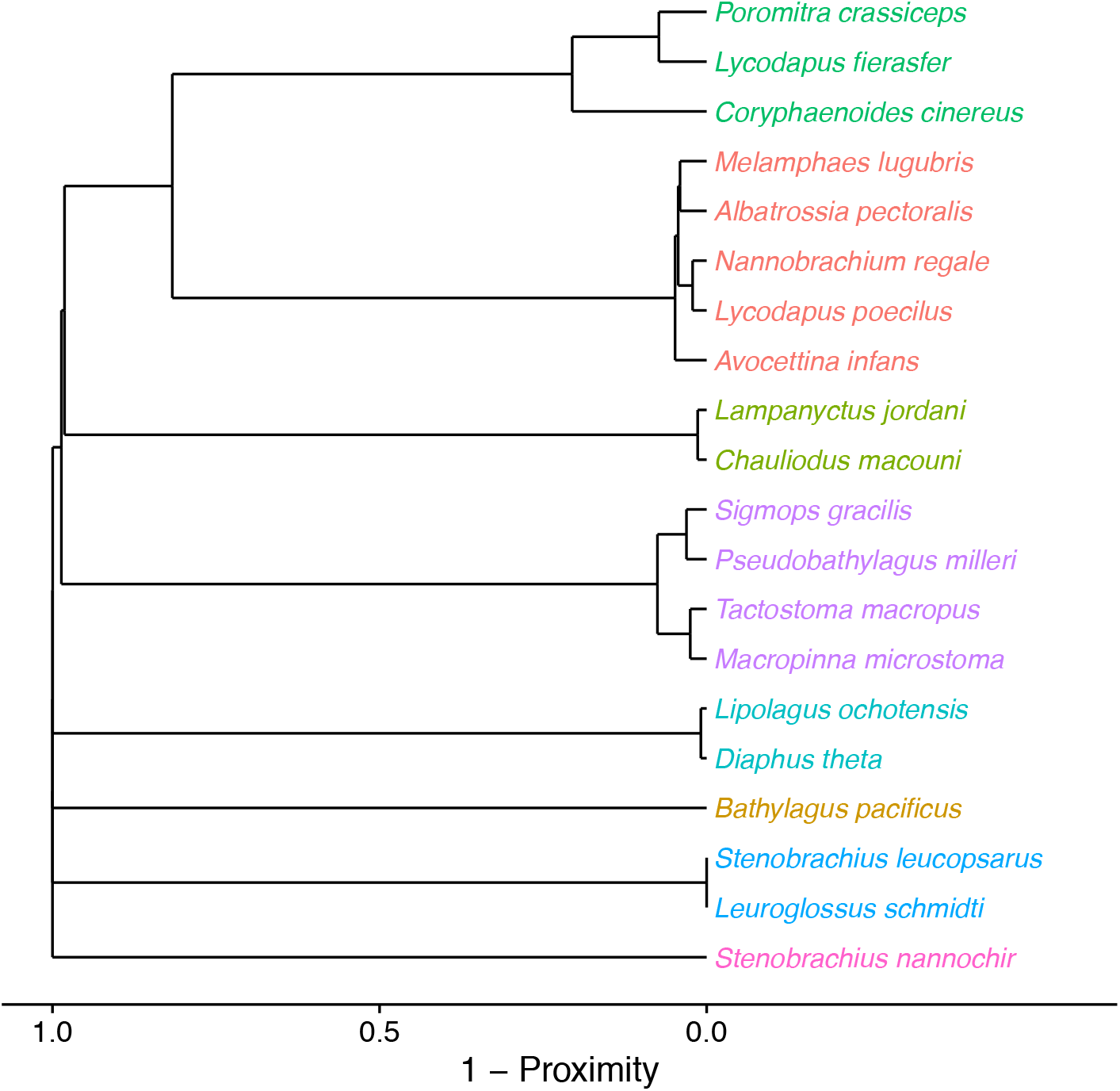
Clustering of trawl survey fish species based on the estimated probability of joint guild membership. The matrix 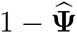 was used as a distance matrix for forming the dendogram. The colored labels reflect guild groupings based on the posterior mode number of guilds, 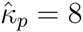

The predicted abundance for each species was calculated as 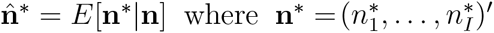 is an observation under the various environmental conditions (Figure 6).

**Figure 6.**
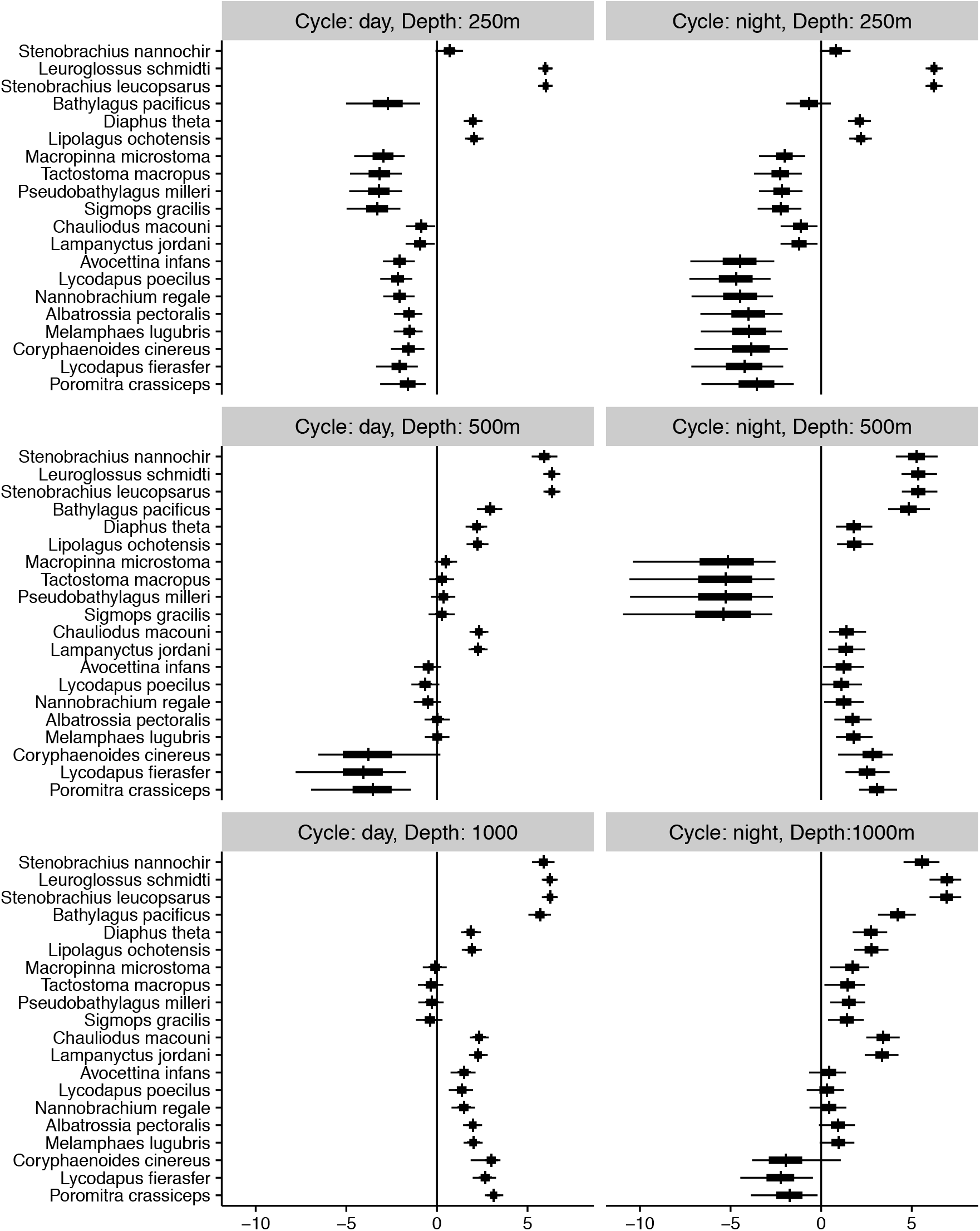
Species-specific predictions of log-abundance for each level of cycle (day or night), and depth (250, 500, or 1000 m).

Results for the γ parameters are presented in Table B.1 of Supplementary Material B along with estimates of the 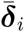 values (Figure B.1). Supplementary Material C provides similar figures and results for the DP-JSDM model using binary occurrence data instead of the observed abundance.

The model profiled a wide range in behavior among species from the two dominant mesopelagic fish families in the Bering Sea, Myctophidae and Bathylagidae. All but one of the 8 guilds described by the model (Figures 5 and C.2) include a single species from one or both of these families, implying that they partition the water column based on a characteristic response to physical factors and foraging requirements.

The accuracy and predictive capability of the model was confirmed by the correct guild assignment of individual species with previously known abundance and depth distribution profiles in the Bering Sea (i.e., bathylagids, *Leuroglossus schmidti* and *Lipolagus ochotensis).* Then by virtue of guild membership, the model described distribution patterns of species for which there is little reported data (i.e., myctophids, *Stenobrachias leucopsarus* and *Diaphus theta*).

For instance, *L. schmidti* and *S. leucopsarus* formed the tightest cluster in both abundance and occurrence dendograms (Figures 5 and C.2). Each is the most abundant species within their respective families in the Bering Sea (Brodeur *et al.,* 1999; Sinclair *et al.*, 1999) and both were highly represented throughout the water column day and night in this study. Guild membership with *L. schmidti* suggests that *S. leucopsarus* shares a similar life history and foraging strategy wherein juveniles and adults have indistinct vertical migration and are stratified in the water column according to age (size) with adults remaining below 240 m (Beamish *et al.*, 1999; Mecklenburg *et al.*, 2002).

The bathylagid *L. ochotensis* and myctophid *D. theta* also form a guild in abundance (Figure 5) along with *Stenobrachias nannochir* in occurrence guilds (Figure C.2). *Lipolagus ochotensis* and *S. nannochir* are among the most abundant mesopelagic species in the Bering Sea (Sinclair *et al.*, 1999; Mecklenburg *et al.*, 2002). Both are size-stratified by depth with adults residing in the deepest layers and especially present between 500–1000 m (Mecklenburg *et al.*, 2002). As a strong vertical migrator, *L. ochotensis* is also abundant between 200–500 m (Sinclair *et al.*, 1999; Mecklenburg *et al.*, 2002). Little is known about *D. theta* from directed catch in the Bering Sea, however guild identity with *S. nannochir* and especially with *L. ochotensis* suggests they share similar patterns of behavior. The model implication that *D. theta* is an age-stratified strong vertical migrator available at upper mesopelagic depths (Figure 6, B.1, and C.3) is supported by observations that it is a primary prey item of Dall’s porpoise (*Phocoenoides dalli*) in the top 250 m of water column (Crawford, 1981).

The best example of the degree of fine detail captured by the model was demonstrated by *Bathylagus pacificus,* a common and abundant species of Bathylagidae that formed its own cluster (Figure 5). Like other members of its family *B. pacificus* demonstrates a bimodal pattern in body size at depth (Peden *et al.*, 1985; Mecklenburg *et al.,* 2002). In our study, juvenile fish were concentrated at mid-layer levels during the day (500 meters) rising to 250 meters at night, while adults concentrate at deeper daytime layers (1000 m) rising to 500 m at night (Sinclair and Stabeno, 2002). This vertical migratory movement is apparent in the log abundance plots (Figure 6; and 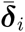 values in Figure B.1) that together with known age distribution suggest *B. pacificus* may form its own guild based on abundances at depth driven by varying foraging requirements of juvenile and adults.

## 5 DISCUSSION

We presented a new methodology for modeling joint species distributions based on Dirichlet process random effects to model species associations through a latent guild structure. Instead of trying to directly parameterize cross-correlation in a species-specific random effect, we used latent membership in an ecological guild. Species belonging to the same guild followed the same response to environmental conditions through random coefficients effects in a GLM-like setting. Unlike simple cross-correlated species random intercepts, the DP-JSDM provides some valuable information on which species belong to guilds together and for the species within a guild, how they respond to the selected environmental conditions together.

A fundamental aspect of mesopelagic ecology is diel vertical migration. The DP-JSDM successfully identified community structure among 20 species of fish from the eastern Bering Sea within this framework. The selected model parameters of depth and light describe realtime clusters of species that move together similarly through the water column on a 24 hour cycle, presumably in relation to foraging. Based on studies conducted in the North Pacific Ocean, the diets of many of these same species collected from different depths match vertical distribution patterns of the variety of copepods and euphausiids that they consume (Beamish *et al.*, 1999).

Although the DP-JSDM model was initially designed to model species association, it has the added benefit that it automatically adjusts to the necessary complexity because the number of guilds is also simultaneously being estimated as well. In the simulation experiment it was demonstrated that if there is little difference between the species in their response to the recorded environmental conditions the DP-JSDM will collapse to one guild, that is, no statistical difference between the species. This reduction in model complexity was noted by Johnson *et al.* (2013b) in reference to spatially clustering abundance trends.

In our description of the model and our examples, we have provided a relatively straightforward demonstration of the model and associated RJMCMC algorithm. However, there are several extensions that would be useful in other ecological settings. Here we did not have repeated observations at each site, so, we could not add an identifiable detection model to the observation process, although, we illustrated that covariates (i.e., trawl duration) could be added as a quasi-detection model as Ver Hoef and Frost (2003) used. However, if multiple observations are available for each site, then a detection process could be added to the observation model. Dorazio and Connor (2014) made use of an N-mixture model and the DP-JSDM could use that as well. Instead of the ZIP model, one could add a another observation model,

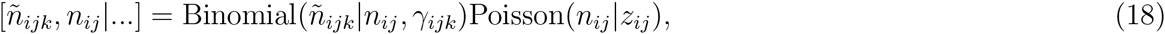

as the observation portion of the model, where *ñ_ij_* is the observed abundance of species *i* at site *j* during survey *k* and *γ_ijk_* is the probability of each of the *n_ij_* individuals being observed. If one marginalizes over the true abundances, the Poisson observation model results,

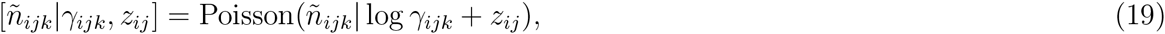

where *E*[*n_ijk_*] = exp{log *γ_ijk_* + *z_ij_*}. The same approach could also be used for occurrence modeling, in which case, it becomes occupancy modeling, that is, for the observed presence 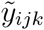, we use the hierarchical observation model,

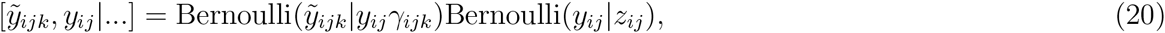

where the probability that 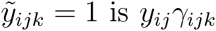. The main point being that the process model does not change in either of these two settings, so, the DP-JSDM can easily be adapted to these situations.

There is also an alteration that can be made when many sites are visited and spatial correlation between sights might also be a consideration. We are not calling this an extension, because spatial correlation can be added without making additions to the basic structure presented. All that needs to be changed to add random spatial effects is to use the basis function approach of Ver Hoef and Jansen (2014), Johnson *et al.* (2013a), or Hefley *et al.* (2016). In a spatial basis function model, the random spatial field is modeled as ***η*** = **H*δ*** where the columns of the matrix **H** contain the spatial basis functions evaluated at each of the modeled sites (rows). Each basis column represents a different frequency. In the notation just presented it should be fairly obvious how the DP-JSDM can be changed to contain spatial correlation, one simply needs to use a basis function matrix for the environmental design matrix. In that case, it might be appropriate to use 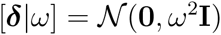 for the DP baseline distribution to match prior specifications that are usually used in spatial analysis. And, of course, one could combine the spatial model with the previously mentioned detection model extensions to form mutivariate spatial models for occupancy and abundance modeling.

## ACKNOWLEDGEMENTS

The authors thank M. B. Hooten and J. L. Laake for a critical reading of the original version of the paper. The findings and conclusions in the paper are those of the authors and do not necessarily represent the views of the National Marine Fisheries Service, NOAA. Reference to trade names does not imply endorsement by the National Marine Fisheries Service, NOAA.

## Footnotes

1 Available from github at: https://github.com/dsjohnson/multAbund. The package can be installed from within an `R` session using the `devtools` package, but users need to be able to compile source code on their platform as the `multAbund` package uses `C++` code in its routines.

## Supplementary Material A: RJMCMC Details

### 1 Prior distributions

Here we describe the details for performing the necessary parameter updates in the RJMCMC algorithm. To facilitate the description the reader should recall we use the following prior distributions in full vector form (where appropriate):

- 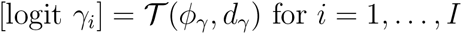
- 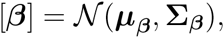
- 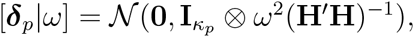
- 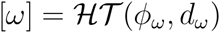
- 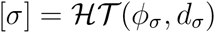
- 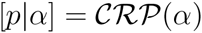
- 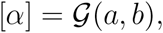

where 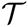 denotes a *t* distribution, 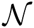 is a (multivariate) normal distribution, 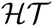 is a half-t distribution, 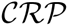 is the Chinese restaurant process, and 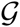 is a gamma distribution. Now, we can describe the Markov Chain Monte Carlo (MCMC) sampler. The sampler is constructed from repeated draws from the full conditional posterior distributions. We use the notation [*x*|·] to represent the conditional distribution of the variable ‘*x*’ given all of the other model components.

### 2 Updating z

We will first describe the updating of **z** for the abundance models. Unfortunately, for the abundance models used in this paper (e.g., [*n_ij_*|*z_ij_*, *γ*] = ZIP or Poisson), the full conditional distribution does not exist in a nice closed form and we suspect this is the case for every abundance model one may want to use. The full conditional distribution required for the update is,

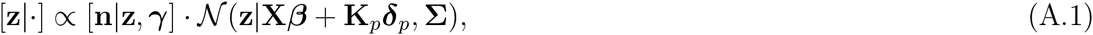

for which a Metropolis-Hastings (MH) step is used with a random walk proposal distribution [**z**\*|**z**] = *N*(**z**, **R***_z_*), where **R***_z_* is a diagonal matrix that is tuned for optimal sampling. In the `R` package `multAbund` we use the adaptive random walk proposal described by Shaby and Wells (2011) that continually adjusts proposal distribution throughout the MCMC run. Once the new **z**\* is drawn, each 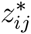 is accepted with probability

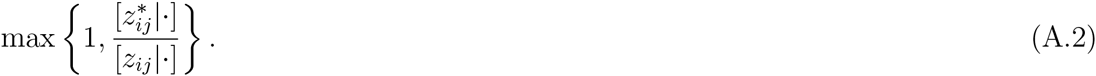

Note, that even though **z**\* is proposed as a vector, the independence of each element implies that each 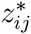 can be accepted or rejected independently.

If one is analyzing occurrence data with a probit link as described in the main text of the paper, then the full conditional distribution,

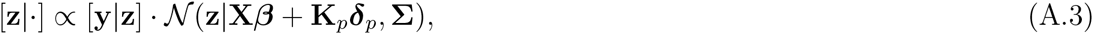

is available in closed form. For each (*i*, *j*), the necessary full conditional distribution is

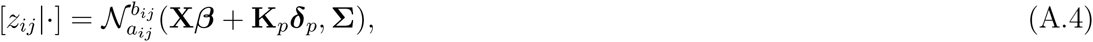

where 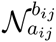 is a truncated normal distribution with lower bound

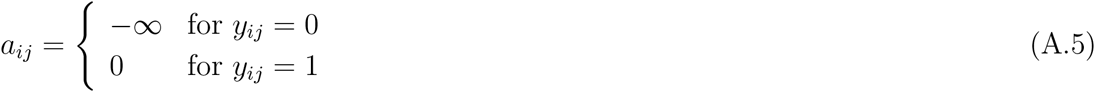

and upper bound

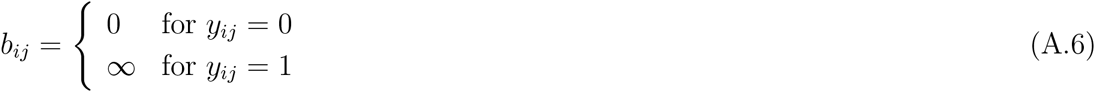

(Albert and Chib, 1993). If another link function is used, then the same procedure as the abundance model updates is used with a MH acceptance step.

### 3 Udating γ

Here, the only model used where γ was present is the ZIP model used in the analysis of the fish survey data. Therefore, we only describe updating of this parameter with respect to the ZIP model with species-specific ZIP parameters, γ*_i_*. The full conditional distribution of logit γ*_i_* is

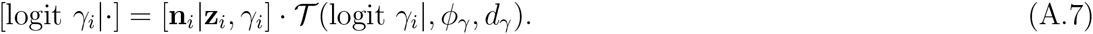

As with the **z** updates, the adaptive random walk MH update 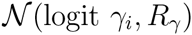 was used were *R*_γ_ is continually adapted through the RJMCMC.

### 4 Updating *β* and *δ_p_*

All of the coefficient vectors in the model have a normal prior distribution, thus the full conditional distributions [*β*|·] and [***δ**_p_*|·] are normal distributions where each is given in Table A.1.

**Table A.1:** Means and variances for sampling of *β* and ***δ**_p_*. Each parameter has a full conditional distribution of the form 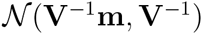.

| Distribution | $\mathbf{V}$ | $\mathbf{m}$ |
| --- | --- | --- |
| $[\beta\vert \cdot]$ | $\mathbf{X}'\Sigma^{-1}\mathbf{X} + \Sigma_{\beta}^{-1}$ | $\mathbf{X}'\Sigma^{-1}(\mathbf{z} - \mathbf{K}\delta_p) + \Sigma_{\beta}^{-1}\mu_{\beta}$ |
| $[\delta_p\vert \cdot]$ | $\mathbf{K}_p'\Sigma^{-1}\mathbf{K}_p + (\mathbf{I}_{\kappa_p} \otimes \Omega)^{-1}$ | $\mathbf{K}_p'\Sigma^{-1}(\mathbf{z} - \mathbf{X}\beta)$ |

### 5 Updating *ω* and *σ*

Using an 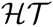 family of priors is not directly conjugate, therefore, a MH step is used here as well. Recall that here we are using **Ω** = *ω*^2^(**H′H**)^−1^ and Σ = *σ*^2^**I**, where *ω* = exp(*ξ*) and *σ* = exp(*θ*). These choices could be easily modified if desired. For *ω*, the full conditional distribution is given by

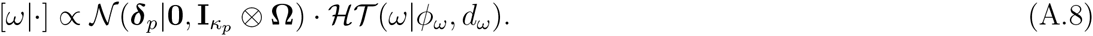

when converting to the log parameterization, we obtain the full conditional for *ξ*,

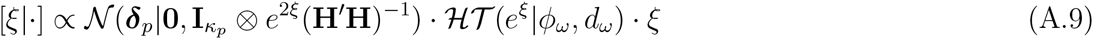

As in the *z* updates, we use a normal random-walk proposal 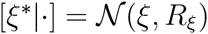, where *R_ξ_* is adaptively tuned throughout the MCMC run in the way as the **z** updates. With regards to *σ*, the *θ* parameter is updated in an identical fashion with the full conditional distribution given by

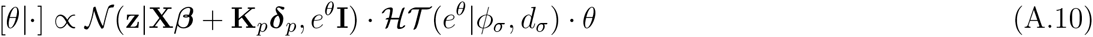

and adaptive random walk proposal distribution 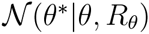.

### 6 Updating *p* and *α*

The update of *p* was described in the main portion of the paper, therefore we omit it here and refer the reader to Section 2.2 for details.

The CRP parameter *α* is updated through an MH step with the previously described adaptive random walk proposal on log a. The full conditional distribution is given by

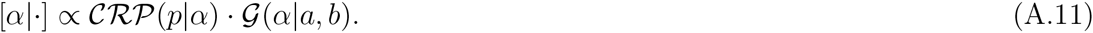

However, as with all of the positive valued parameters, we choose to reparameterize to the log scale to make use of the adaptive random walk proposal distribution. So, the full conditional distribution for log *α* is

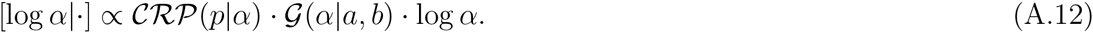

The same adaptive procedure was used with an MH acceptance step to sample the full conditional distribution.

### Acknowledgments

The findings and conclusions in the paper are those of the authors and do not necessarily represent the views of the National Marine Fisheries Service, NOAA. Reference to trade names does not imply endorsement by the National Marine Fisheries Service, NOAA.

## Supplementary Material B: Additional results for fish survey abundance model

**Table B.1:** Results for species-specific Zero-inflated Poisson (ZIP) mixture parameters, γ*_i_*. The ‘Estimate’ column is the posterior mode estimate and the ‘CI’ columns are the upper and lower 0.95 highest probability density interval values. The mixture probabilities represent the probability that a given species is unavailable for surveying in a particular survey.

|  | Estimate | Lower CI | Upper CI |
| --- | --- | --- | --- |
| *Albatrossia pectoralis* | 0.20 | 0.03 | 0.44 |
| *Avocettina infans* | 0.52 | 0.27 | 0.74 |
| *Bathylagus pacificus* | 0.04 | 0.00 | 0.20 |
| *Chauliodus macouni* | 0.03 | 0.00 | 0.17 |
| *Coryphaenoides cinereus* | 0.14 | 0.00 | 0.46 |
| *Diaphus theta* | 0.18 | 0.04 | 0.33 |
| *Lampanyctus jordani* | 0.08 | 0.00 | 0.24 |
| *Leuroglossus schmidtii* | 0.01 | 0.00 | 0.07 |
| *Lipolagus ochotensis* | 0.13 | 0.02 | 0.28 |
| *Lycodapus fierasfer* | 0.43 | 0.20 | 0.69 |
| *Lycodapus poecilus* | 0.58 | 0.35 | 0.79 |
| *Macropinna microstoma* | 0.07 | 0.00 | 0.36 |
| *Melamphaes lugubris* | 0.18 | 0.00 | 0.40 |
| *Nannobrachium regale* | 0.54 | 0.28 | 0.75 |
| *Poromitra crassiceps* | 0.03 | 0.00 | 0.28 |
| *Pseudobathylagus milleri* | 0.28 | 0.00 | 0.56 |
| *Sigmops gracilis* | 0.37 | 0.03 | 0.66 |
| *Stenobrachius leucopsarus* | 0.01 | 0.00 | 0.07 |
| *Stenobrachius nannochir* | 0.04 | 0.00 | 0.15 |
| *Tactostoma macropus* | 0.32 | 0.00 | 0.57 |

**Figure B.1:**
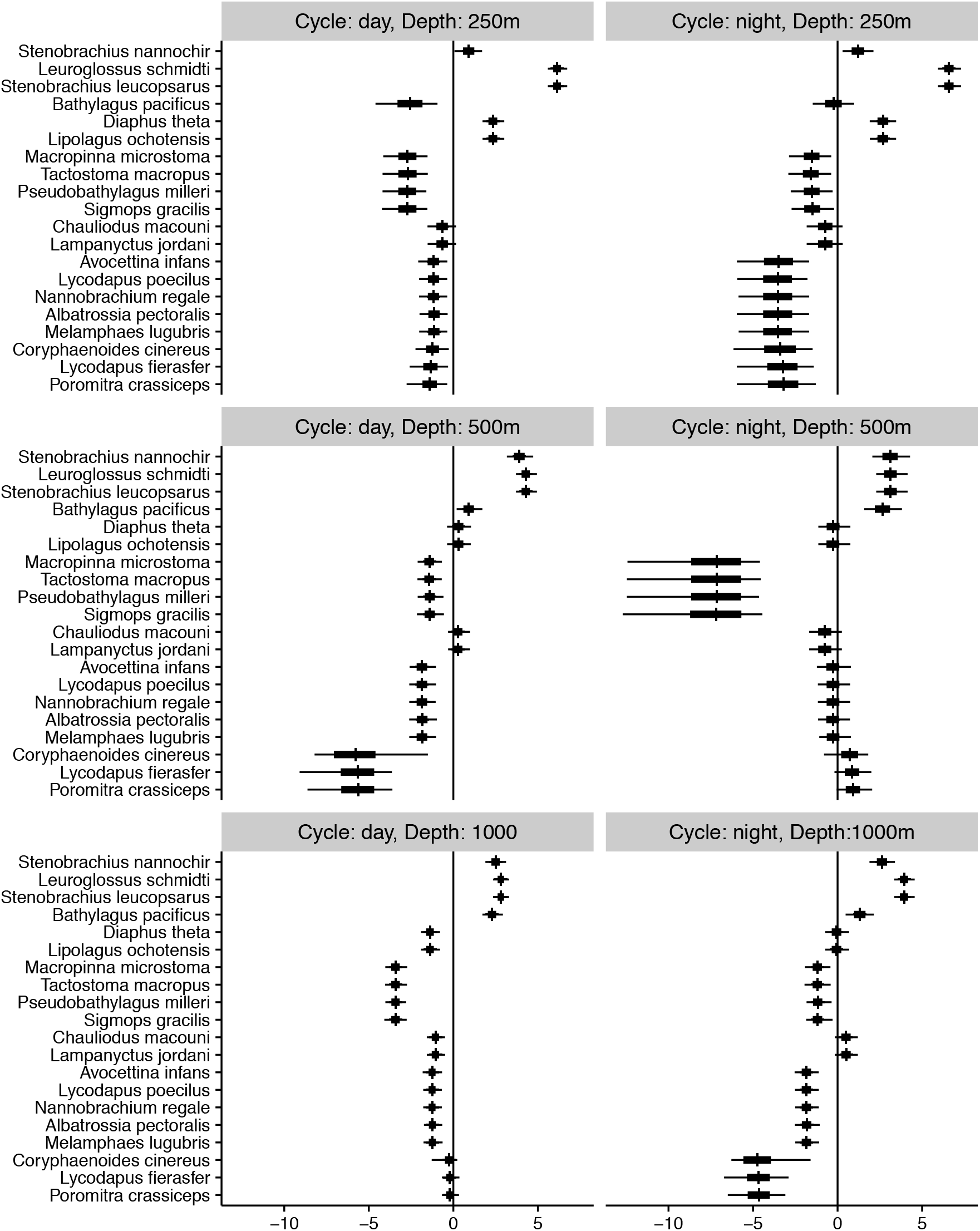
Species-specific *δ* estimates, 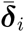, for each level of cycle (day or night), and depth (250, 500, or 1000 m).

## Supplementary Material C: Mesopelagic fish survey occurrence modeling

**Figure C.1:**
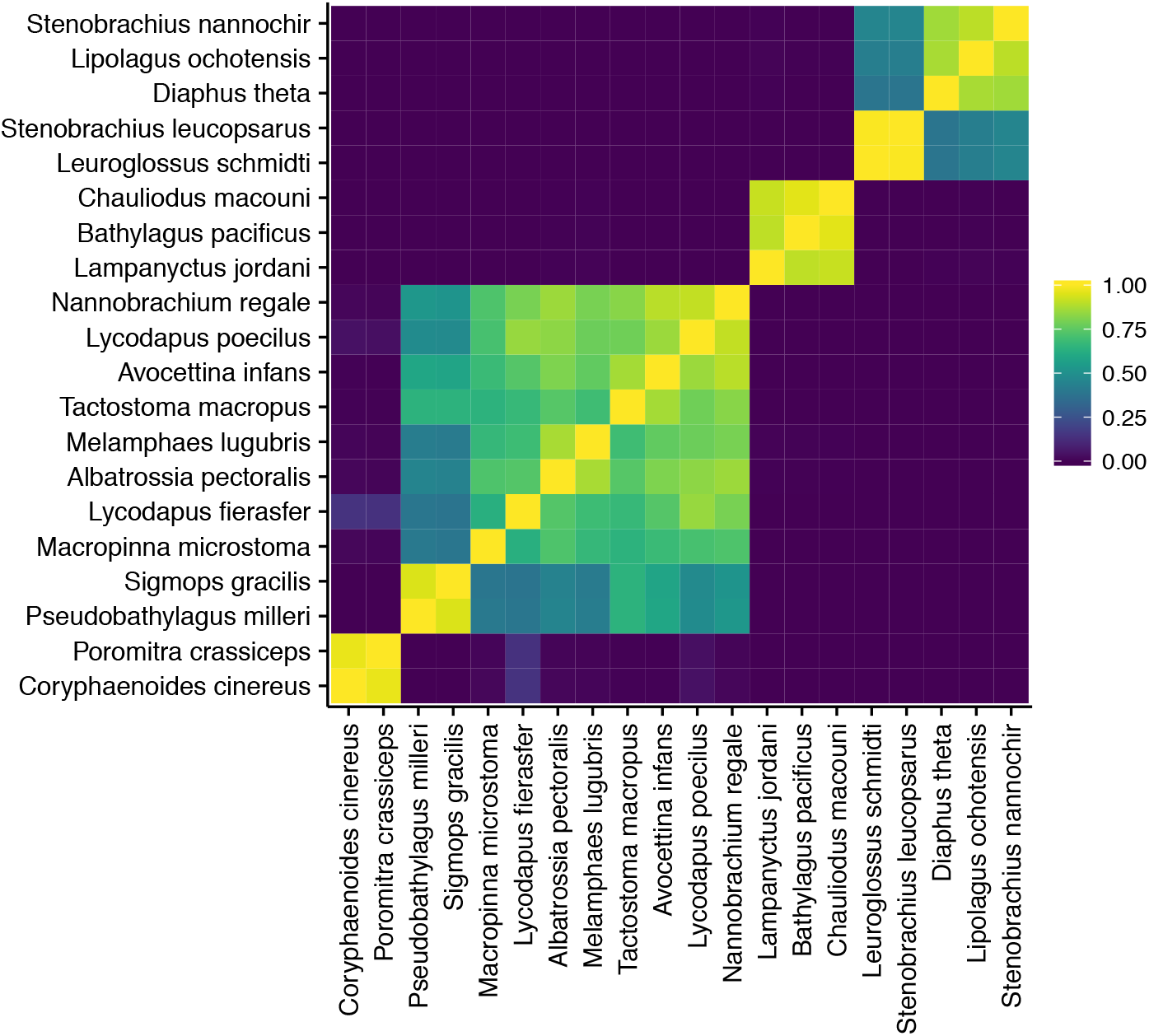
Estimated probability of joint guild membership for each of the fish species in the trawl survey with respect to occurence.

**Figure C.2:**
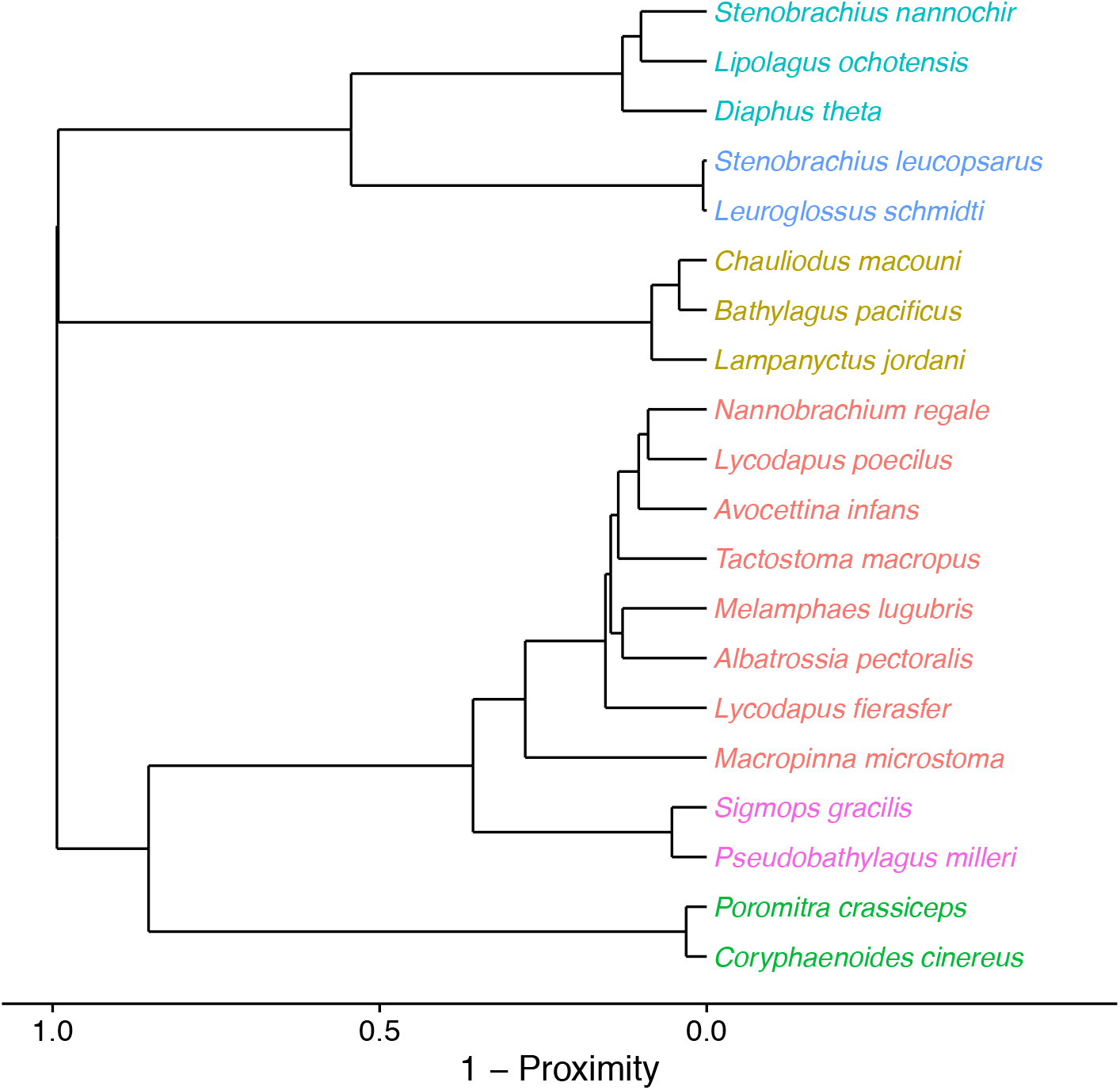
Clustering of trawl survey fish species occurrence based on the estimated probability of joint guild membership.

**Figure C.3:**
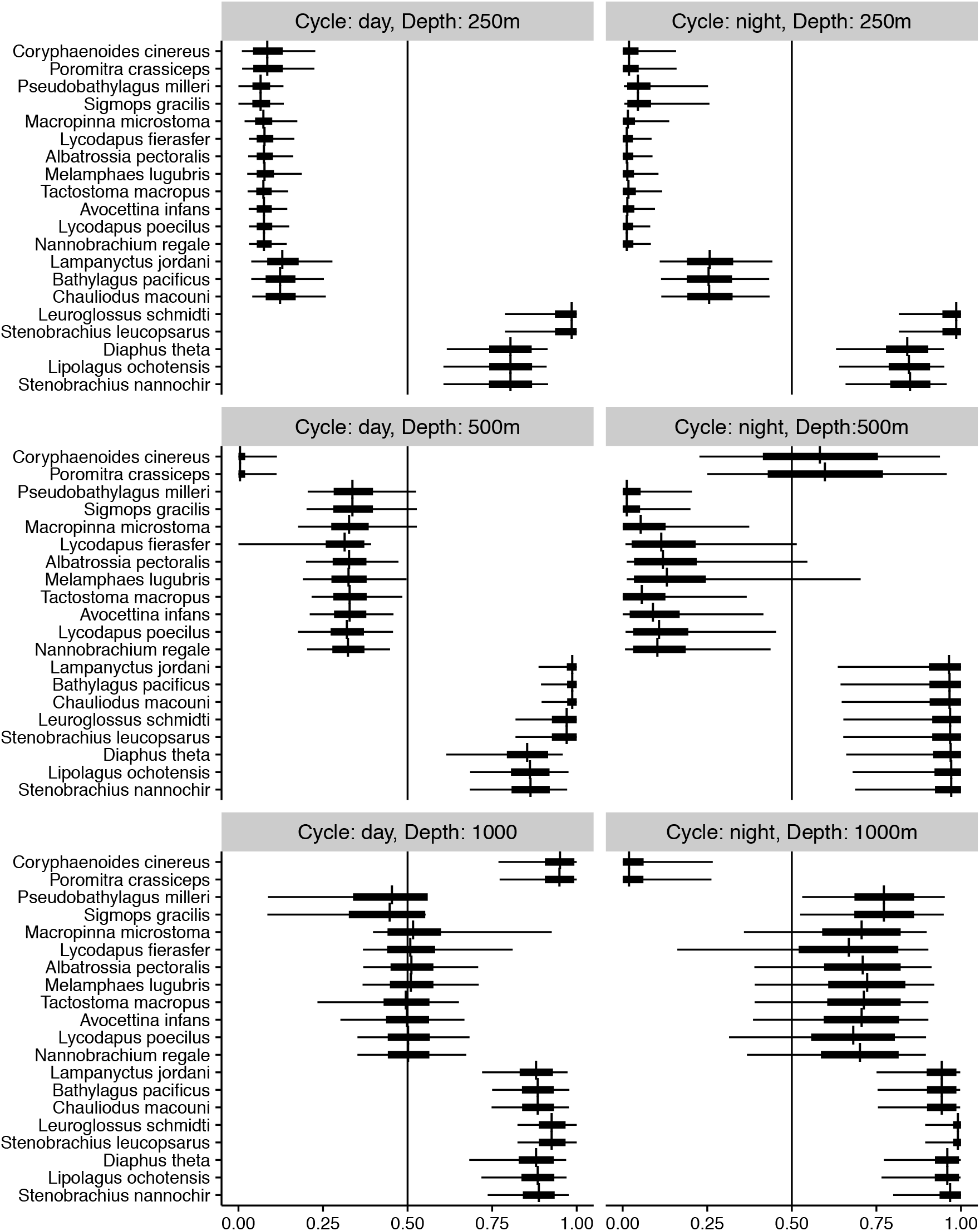
Species-specific predictions of occurrence for each level of cycle (day or night), and depth (250, 500, or 1000 m).

